# Alpha-XIC: a deep neural network for scoring the coelution of peak groups improves peptide identification by data-independent acquisition mass spectrometry

**DOI:** 10.1101/2021.04.20.440630

**Authors:** Jian Song, Changbin Yu

## Abstract

**Motivation:** The peptide-centric identification methodologies of data-independent acquisition (DIA) data mainly rely on scores for the mass spectrometric signals of targeted peptides. Among these scores, the coelution scores of peak groups constructed by the chromatograms of peptide fragment ions have a significant influence on the identification. Most of the existing coelution scores are achieved by artificially designing some functions in terms of the shape similarity, retention time shift of peak groups. However, these scores cannot characterize the coelution robustly when the peak group is in the circumstance of interference.

**Results:** On the basis that the neural network is more powerful to learn the implicit features of data robustly from a large number of samples, and thus minimizing the influence of data noise, in this work, we propose Alpha-XIC, a neural network-based model to score the coelution. By learning the characteristics of the coelution of peak groups derived from identified peptides, Alpha-XIC is capable of reporting robust coelution scores even for peak groups with interference. With this score appending to initial scores generated by the accompanying identification engine, the ensuing statistical validation tool can update the identification result and recover the misidentified peptides. In our evaluation of the HeLa dataset with gradient lengths ranging from 0.5h to 2h, Alpha-XIC delivered 16.7% ~ 49.1% improvements in the number of identified precursors at 1% FDR. Furthermore, Alpha-XIC was tested on LFQbench, a mixed-species dataset with known ratios, and increased the number of peptides and proteins fell within valid ratios by up to 16.6% and 13.8%, respectively, compared to the initial identification.

**Availability and Implementation:** Source code are available at www.github.com/YuAirLab/Alpha-XIC.

## 1 Introduction

Data-independent acquisition (DIA) mass spectrometry (MS) has been widely used in proteomics due to its unbiased and systematic measurement of precursors and fragment ions compared to data dependent acquisition (DDA), which improves peptide detection and quantification in the analyses of complex biological samples (Ludwig *et al*., 2018). The prevalent strategy of peptide identification for DIA data is currently the peptide-centric matching approach (Duncan *et al*., 2010; Zhang *et al*., 2020). For each targeted peptide, this approach extracts chromatograms (also referred to as traces which are continuous in retention time and intensity) of a certain number of most intensive fragment ions and assembles them into a peak group followed by scoring, and finally performs statistical validation based on the scores using target-decoy methods (Röst *et al*., 2014). In general, it is necessary to score the peak group in terms of the mass accuracy, the intensity similarity between the experimental relative intensities and the relative intensities stored in the spectral library, the deviation between the expected and measured retention time, and the coelution of the traces (Reiter *et al*., 2011). Since peak groups are a complete two-dimensional record of the fragment ions and the coelution reflects the consistency of the fragment ions in dimensions of retention time and intensity, how to score the coelution of peak groups has a great impact on DIA identification.

Different DIA identification engines have different strategies to score the coelution. Skyline (MacLean *et al*., 2010) used dot product to represent the coelution of peak groups, which is over-simplistic, especially when the peak group is disturbed. To score coelution comprehensively and robustly, OpenSWATH (Röst *et al*., 2014) calculated five scores to evaluate the coelution including the dot product, the cross-correlation and the global retention time shift both weighted or non-weighted by the relative intensities of fragment ions. Alternatively, DIANN (Demichev *et al*., 2020) selected a ‘best’ fragment trace that was considered the one least likely to be affected by interference and scored the coelution using Pearson correlation between each trace with the ‘best’ fragment trace. To make sure that the coelution score was not dominated by a single highly scored trace, Avant-garde tool (Jacome *et al*., 2020) removed the trace with the highest dot product and calculated a second mean of the remaining dot products. Although the above coelution scores have attempted to reduce the effects of interference as much as possible, the complex and varied forms of interference, such as signal loss, jagging or convolution, still make them susceptible and far from robust.

Deep learning, or deep neural network (LeCun *et al*., 2015), as a method to learn data inherent characteristics powerfully from a large amount of samples, and thus minimizing the influence of data noise, is suitable for the robust coelution scoring problem. In this paper, we present Alpha-XIC, a neural network-based model to score the coelution of peak groups. This model is trained as a classifier on part of peak groups determined by an accompanying DIA engine and then expanded to score coelution for all candidate peak groups. After appending the output of the model as an additional score to initial scores, the statistical validation tool can update the identification result. Our preliminary experiments indicate that Alpha-XIC can score the coelution of peak groups robustly and thus significantly improving the identification of DIA data.

## 2 Methods and material

### 2.1 Workflow

There are a few factors that affect the appearance of peak groups, such as liquid chromatography (LC) gradient lengths, cycle strategies of DIA experiments and noise baselines of mass spectrometers (Ludwig *et al*., 2018). In order to make Alpha-XIC more specific to the peak groups that need to be scored, it is not designed to be a model that is trained once and used for each DIA file. Instead, Alpha-XIC is trained on the peak groups derived from the being analyzed DIA data. From this point of view, Alpha-XIC depends on an DIA engine to perform an initial identification to offer training samples. In other words, Alpha-XIC can be considered as a plug-in of the DIA engine. Since OpenSWATH is convenient to add custom score, it is adopted as the matching engine for Alpha-XIC in this study. The workflow of Alpha-XIC was plotted in Fig. 1 and described in phases as follows.

1. Initial identification by OpenSWATH and PyProphet (Teleman *et al*., 2015). With the help of the spectral library and the preprocess of retention time alignment, OpenSWATH extracted the fragment ion traces, determined and scored the peak group for each target or decoy peptide. Then, PyProphet, a statistical validation tool, was employed to calculate the false discovery rate (FDR) by the target-decoy method. At the end of this phase, candidate peak groups were obtained as well as their assigned FDR.
2. Training and utilization of Alpha-XIC. We selected the peak groups from the previous phase within 1% FDR as positive samples. To avoid the potential class imbalance problem, an identical number of decoy peak groups were picked out randomly from candidate peak groups as negative samples. Then, Alpha-XIC was trained as a binary classifier on these positive and negative samples. After the training was completed, all the candidate peak groups were scored by Alpha-XIC. The resulting positive classification probabilities were used as the additional scores and appended to scores reported by OpenSWATH in phase 1.
3. Update the statistical validation. PyProphet was relaunched after obtaining the new scores by Alpha-XIC to update the statistical validation result. As the FDR for the candidate peak groups had changed after update, the phases 2 and 3 were repeated until either the number of identified peptide at 1% FDR stopped increasing or the maximum number of iterations was reached (5 by default).

**Fig. 1.**
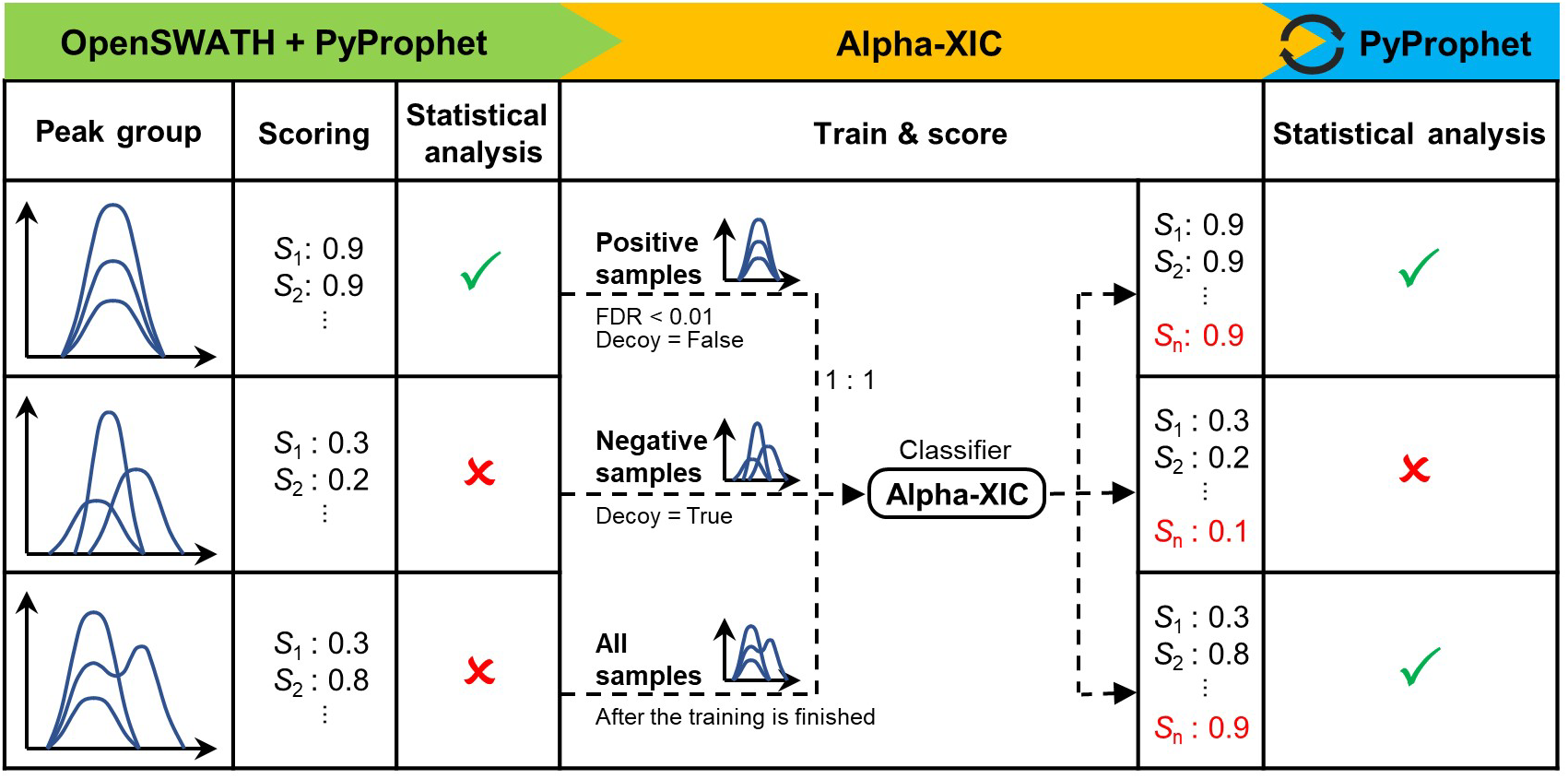
The workflow of Alpha-XIC. From left to right, the workflow of Alpha-XIC includes the initial identification by OpenSWATH and PyProphet, the train and score phase of Alpha-XIC and the update phase of the statistical analysis. The last two phases can be repeated until the stop condition is met. In this figure, the peptide corresponding to the third peak group which has interference is recovered after adding the coelution score by Alpha-XIC.

### 2.2 Model

Under different LC-MS conditions and DIA configurations, peak groups have different peak widths and the number of data points (Ludwig *et al*., 2018). In order to process these diverse peak groups, it is necessary to perform preprocessing to uniform the format of peak groups. The top panel of Fig. 2 plotted the details of preprocessing. For each peak group, Alpha-XIC extracted the traces within the boundaries which were determined by OpenSWATH. As some peak groups were used in both training and scoring phases, we shuffled the order of traces of the peak group randomly in the training phase to avoid possible overfitting. Next, each trace was normalized in intensity and interpolated to a fixed dimension 64. Then, the peak group was smoothed using a Savitzky-Golay algorithm (11 wide and third order) commonly used for spectrometric data suggested by Marc *et al*., 2008 to filter noise.

**Fig. 2.**
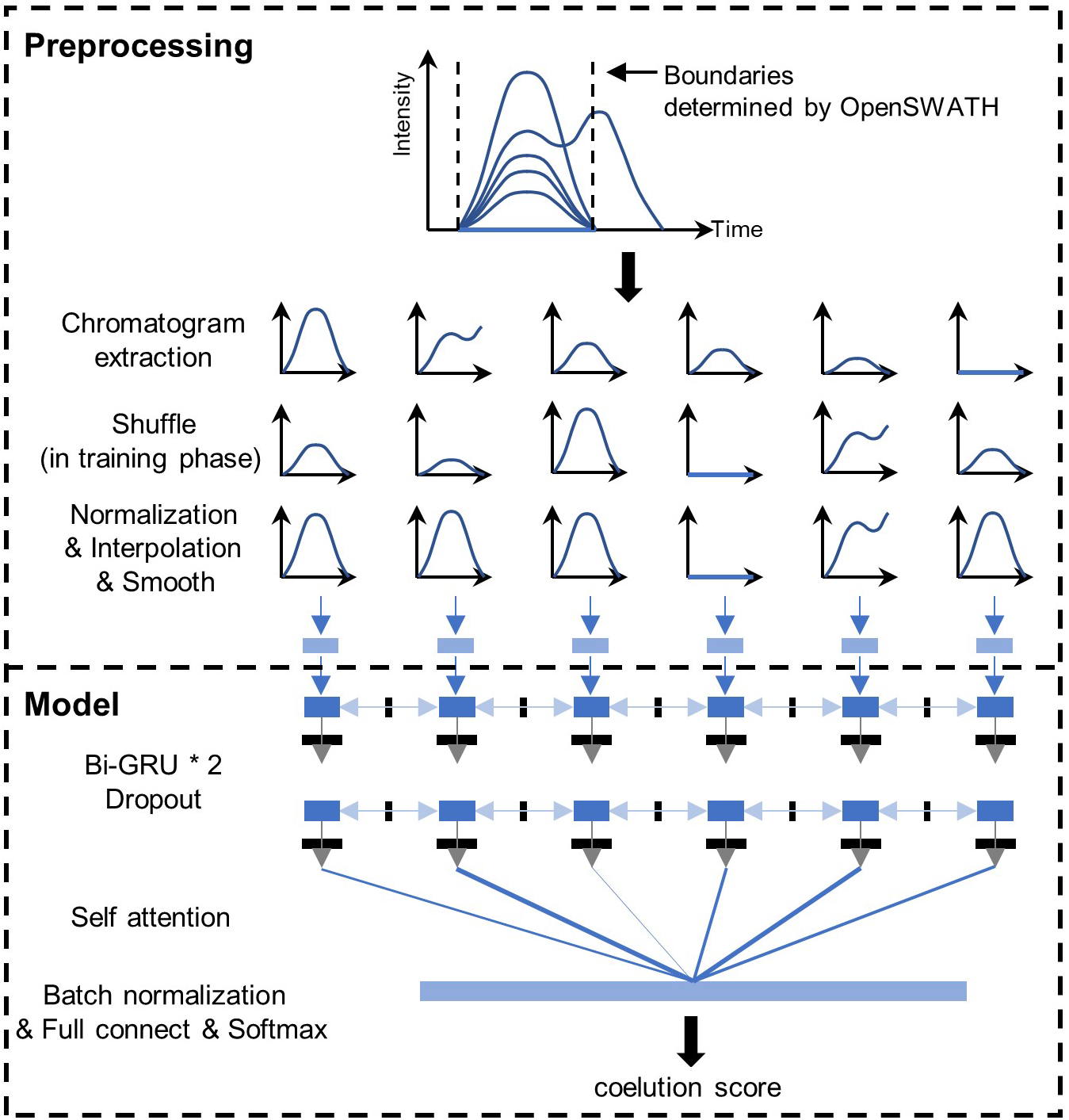
The preprocessing of input peak groups and the model construction of Alpha-XIC.

The bottom panel of Fig. 2 displayed the model components of Alpha-XIC. In order to adapt Alpha-XIC to peak groups with different numbers of traces, RNN (recurrent neural network) was used to analyze the peak group by the means of trace series. Specifically, two layers of bi-directional gated recurrent memory units (Bi-GRU, a type of RNN, Chung *et al*., 2014) with hidden size 64 and dropout (Srivastava *et al*., 2014) 0.5 were selected to process the input of traces. To concatenate the Bi-GRU outputs, a self-attention layer (Lin *et al*., 2017) was used. In this way, the peak group was converted to a vector of dimension 128. Then, this vector was converted to dimension 2 by a batch normalization layer (Ioffe *et al*., 2015), a fully connected layer and a softmax layer in order. At last, the value represented the positive class probability was output as the score of coelution of the peak group. Note that we experimented with different interpolation dimension and output size of Bi-GRU, however, this did not deliver improvements in performance.

PyTorch (v. 1.1.0, https://pytorch.org) was used to implement and train Alpha-XIC. We used the Adam optimizer with an initial learning rate of 0.001 and 128 samples per batch. The loss function in training was cross entropy loss. Particularly, we found that after one training epoch in each iteration for all test files, the loss value stopped falling (data not shown). We suspected that, on the one hand, tens of thousands of training samples in each training were sufficient for Alpha-XIC, on the other hand, it may be relatively effortless for the model to learn coelution characteristics from peak groups. Thus, we fixed the training epoch to one, which was also helpful for avoiding the potential overfitting.

### 2.3 Datasets

The purpose of Alpha-XIC is to score coelution of peak groups robustly and improve DIA identification. In this study, the following public datasets that have been specifically created for testing DIA software were used to evaluate and benchmark Alpha-XIC.

#### HeLa dataset

This DIA dataset is a HeLa whole-proteome tryptic digestion and acquired on a nanoflow liquid chromatography coupled to a Q Exactive HF mass spectrometer with different gradient lengths (Bruderer *et al*., 2017). Here, four RAW files with gradient 0.5h, 1h, 1.5h, 2h (referred as HeLa-0.5h, HeLa-1h, HeLa-1.5h, HeLa-2h) were selected to cover common experimental setups of gradients. All these four files were collected with collision energy 27.5 and with DIA configuration comprising one MS1 survey scan (using a resolution of 120,000, AGC target of 3e6, maximum fill time of 60ms and *m/z* range of 350-1650) followed by a few consecutive isolated MS2 windows of variable width (using a resolution of 30,000, AGC target of 3e6, injection time of auto and minimum *m/z* of 200). The number of variable isolation windows for HeLa-0.5, HeLa-1h, HeLa-1.5h and HeLa-2h were set to 21, 26, 37, 30, respectively.

#### LFQbench dataset

This dataset contained six SWATH (Gillet *et al*., 2012) files and was acquired on a TripleTOF 6600 with 64 variable isolation windows (Navarro *et al*., 2016). Six files were divided into sample A and sample B, each of sample had three replicates. Sample A consisted of mixed tryptic digestion of human, yeast and *Escherichia coli* proteins in 65%, 15% and 20%, respectively. Sample B consisted of proteins same as sample A but mixed in 65% (human), 30% (yeast) and 5% (*E. coli*). As a result, the expected peptide and protein ratios (A/B) are: 1:1 for human, 1:2 for yeast and 4:1 for *E. coli*. Both .raw files of HeLa dataset and .wiff files of LFQbench dataset were converted to mzML format by msconvert.exe from the ProteoWizard package (version 3.0.19133) with 32-bit precision and zlib compression.

For HeLa dataset, the pan-human mass spectrometry library (PHL, Rosenberger *et al*., 2014) was selected as the spectral library. For LFQbench dataset, as the original library provided by LFQbench project was not complete (Navarro *et al*., 2016), we replaced the human peptides in the original library with PHL library. In addition, the CiRT peptides (Parker *et al*., 2015) were used to perform the retention time alignment by OpenSWATH for both two dataset. Recommended by Demichev *et al*., 2020, the parameter ‘rt_extraction_window’ of OpenSWATH should be proportional to the gradient length. Hence, it was set to 600s, 1200s, 1800s, 2400s for HeLa-0.5h, HeLa-1h, HeLa-1.5h, and HeLa-2h, respectively, and 2400s for six LFQbench files. The other options for OpenSWATH (version 2.4.0) included: ‘-batchSize 1000 -readOptions cacheWorkingInMemory -mz_extraction_window 20 -ppm -mz_correction_function quadratic_regression_delta_ppm -min_rsq 0.8’. PyProphet (v.0.24.1) used the default parameters.

## 3 Results

### 3.1 Case comparisons for different interference

As OpenSWATH can report all of its scores for each candidate peak group, here, we compared the robustness of the coelution score by Alpha-XIC with the scores by OpenSWATH using examples of peak groups with interference. As mentioned in above, multiple scores in two dimensions of the retention time shift and shape similarity are calculated to characterize the coelution of peak groups by OpenSWATH. In detail, the coelution scores by OpenSWATH include (Reiter *et al*., 2011; Röst *et al*., 2014):

**dotprod**: It is computed by averaging the dot product values of pair-wise traces of peak groups.
**xcorr**: To achieve more global characteristics of coelution than a simple comparison of apex time shift, OpenSWATH constructs an upper triangular matrix containing the delay which maximizes the cross-correlation of pair-wise traces. *Xcorr* score is the mean plus the standard deviation of the delays in the matrix.
**xcorr_weight**: Similar to *xcorr*, but using the intensity weights to aggregate the delays rather than the mean.
**shape**: Similar to *xcorr*, OpenSWATH constructs an upper triangular matrix containing the maximum cross-correlation value of pair-wise traces. *Shape* score is the mean of the cross-correlation values in this matrix.
**shape_weight**: Similar to *shape*, but using the intensity weights to assemble the cross-correlation values.

The scores of *dotpord, xcorr, xcorr_weight, shape* and *shape _weight* are 1,0,0,1 and 1, respectively, on peak groups with ideal coelution. When coelution is not perfect due to interference, such as signal loss, jagging or convolution, starting point shift, apex point shift and end point shift, the above five scores will deviate more or less from the ideal values. Unlike these scores, Alpha-XIC quantifies coelution by learning from the peak group itself, rather than by manually designing functions, which makes it possible to minimize the impact of interference for coelution scoring.

In order to compare the robustness of Alpha-XIC score with the above scores under different interference, four peak groups of peptides identified exclusively by Alpha-XIC and verified manually in HeLa-1h were picked out to represent different coelution interference. Fig. 3 plotted the peak groups as well as precursor ion chromatograms and their corresponding coelution scores by OpenSWATH and Alpha-XIC. As can be observed that, a trace (y9+, red color) in precursor ‘ITELFAVALPQLLAK_2’ was disturbed which led to the incoordination of its start and apex points against to the global start and apex points of the peak group. As a result, the *dotprod* was low to 0.34. For the precursor ‘ALLNHLDVGVGR_3’, a trace (y5+, green color) had interference and a trace (y10++, blue color) was missing such that all the coelution scores by OpenSWATH (*dotprod*: 0.35, *xcorr*: 6.95, *xcorr_weight:* 5.97, *shape:* 0.65, *shape_weight:* 0.33) had remarkable deviations compared to idea values. Similarly, two traces (y12+, blue color; y11+, brown color) were disturbed in precursor ‘PVAPSGTALSTTSSK_2’ and thus, particularly, the *dotprod* score was as low as 0.29. The peak group signal of ‘GIVGVENVAELKK_3’ was the most complex and resulted in dramatic deviations for all the coelution scores by OpenSWATH (*dotprod:* 0.36, *xcorr:* 8.29, *xcorr_weight:* 6.68, *shape:* 0.40, *shape_weight:* 0.15). Nevertheless, Alpha-XIC scored consistently these four peak groups with high values (1.00, 0.99, 1.00, 0.97, respectively) which made the corresponding peptides recovered even at a cutoff of 1% FDR. In the circumstance that the coelution of the peak group was really far from the ideal as exemplified by the last case, Alpha-XIC still reported a high score 0.97, we speculated that this may be this peak group conformed to the latent characteristics of coelution learned by the network in the training phase or just was similar to one certain positive sample.

**Fig. 3.**
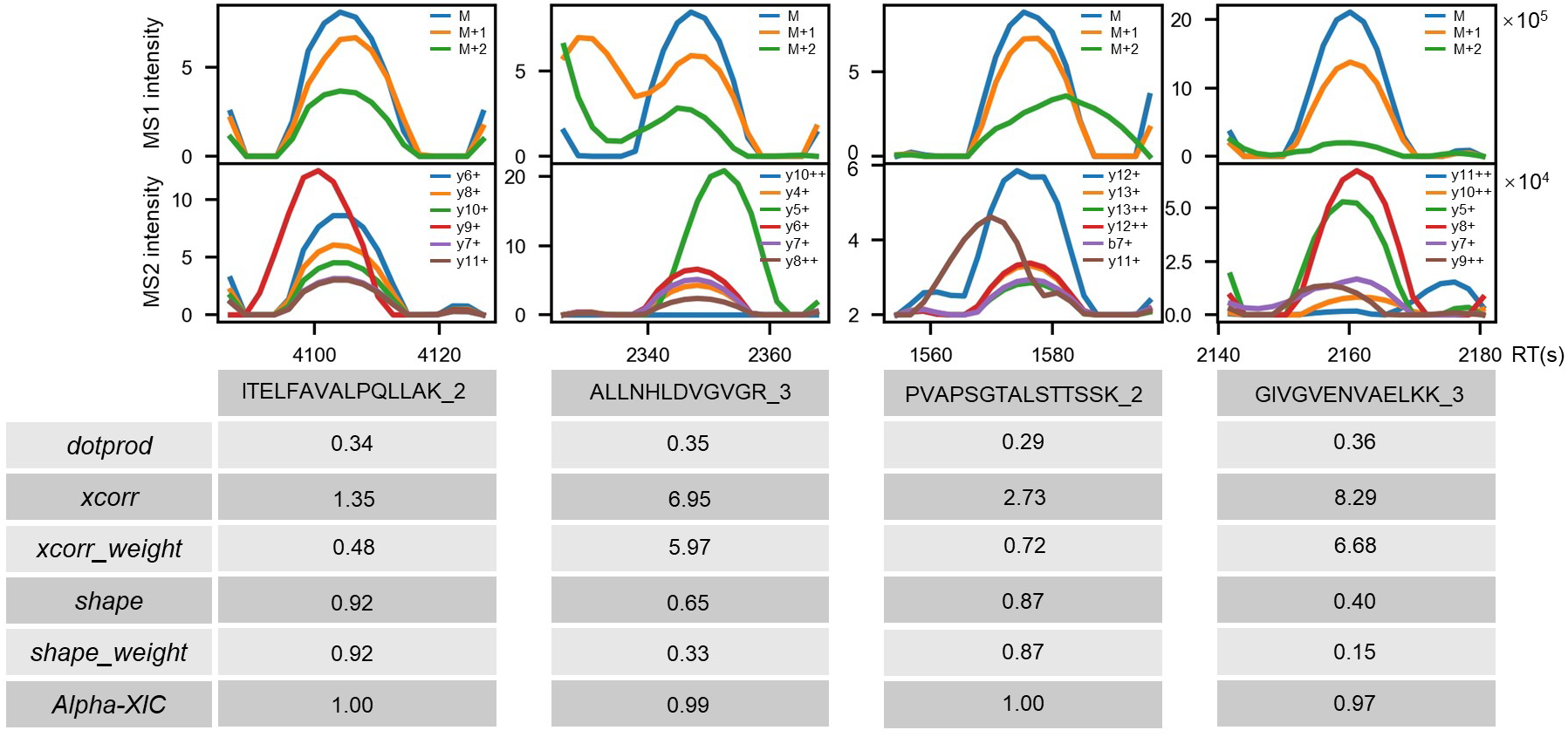
Comparisons of different coelution scores under different interference. The detailed calculation of the coelution scores *‘dotprod’, ‘xcorr’, ‘xcorr_weight’, ‘shape’* and *‘shape_weight* by OpenSWATH is described in Results part. *Alpha-XIC* means the coelution score by Alpha-XIC.

From these case comparisons, it was implied that Alpha-XIC was capable of generating coelution scores robustly.

### 3.2 Performance evaluation on HeLa dataset

To evaluate the performance for DIA identification by Alpha-XIC, OpenSWATH with or without Alpha-XIC were performed to HeLa dataset whose gradient lengths range from 0.5h to 2h in steps of 0.5h. The number of identified precursors was plotted against the FDR in Fig. 4. As can be seen, Alpha-XIC delivered substantial identification improvements than OpenSWATH. At 1% FDR, 49.1%, 27.9%, 24.5% and 16.7% improvements were achieved for HeLa-0.5h, HeLa-1h, HeLa-1.5h and HeLa-2h, respectively.

**Fig. 4.**
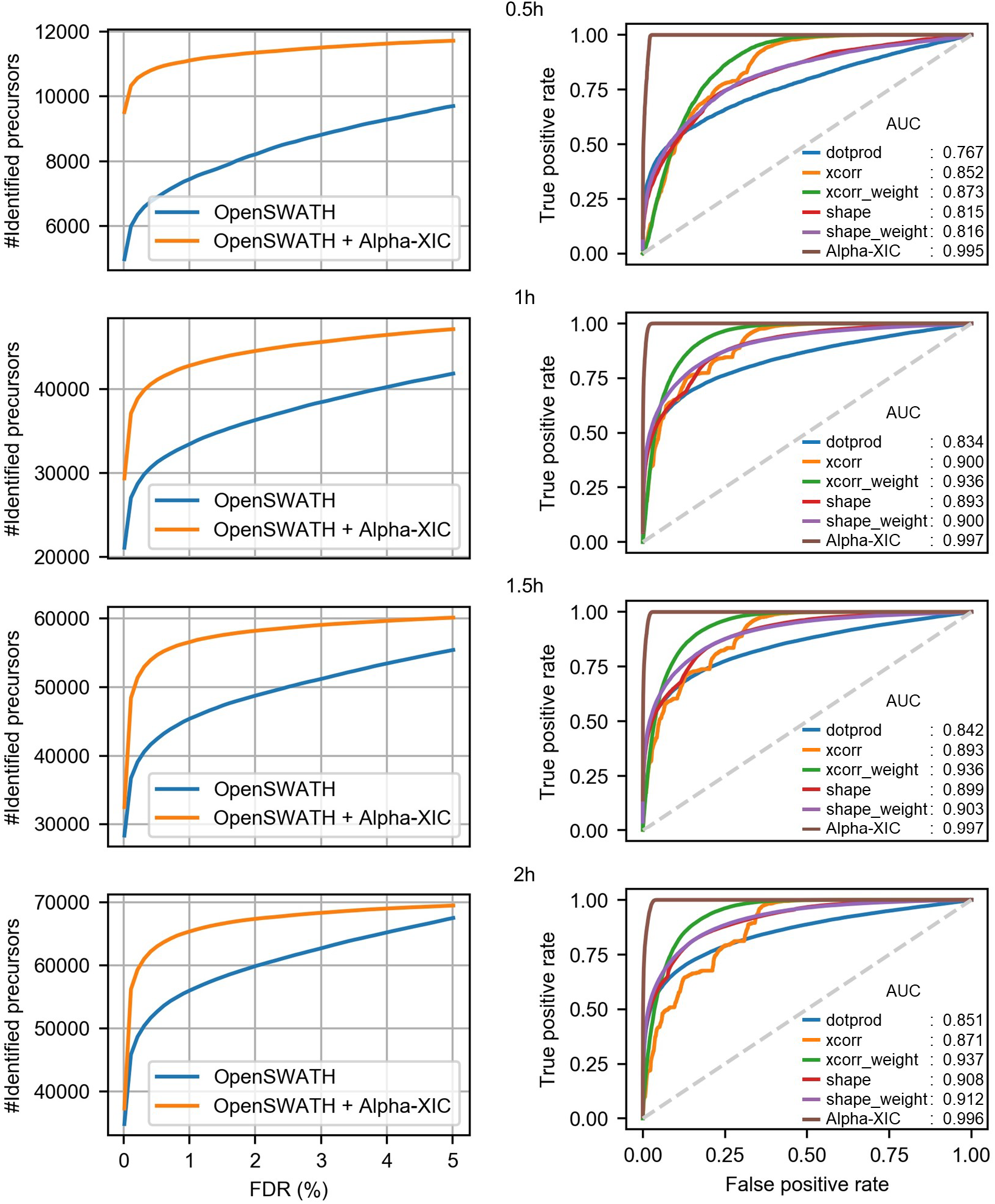
Performance evaluation of Alpha-XIC on HeLa dataset. The gradient lengths of HeLa dataset cover 0.5h, 1h, 1h and 2h. Left panel: the identified precursor numbers against the FDR by OpenSWATH with or without Alpha-XIC. Right panel: ROC plots for each coelution score.

In order to further examine the discriminant ability of the coelution scores by OpenSWATH and Alpha-XIC, separation experiments (Reiter *et al*., 2011) between true and false peak groups were carried out for each individual score. After each HeLa file identified by OpenSWATH combined with Alpha-XIC, we selected the peak groups of the identified target peptides (at 1% FDR) as the positive class, and the peak groups corresponding to the decoy peptides with an equal number by randomly sampling as the negative class. Based on the positive and negative class, we plotted the receiver-operating characteristic (ROC) curve for each individual coelution score. As detailed in the right panel of Fig. 4, the coelution score by Alpha-XIC had the largest area under the curve (AUC) than the coelution scores by OpenSWATH for each HeLa file.

Overall, Alpha-XIC yielded more discriminative coelution score and improved the identification significantly for the DIA data.

### 3.3 Performance evaluation on LFQbench dataset

To assess whether the increase in identification resulting from Alpha-XIC was genuine peptides, we performed the test of Alpha-XIC on LFQbench dataset which had known ratios of mixed species. The LFQbench dataset was identified by OpenSWATH with or without Alpha-XIC, and the quantitative information of the identified precursors under 1% FDR of six files (two types of samples and both with triplicates) was obtained. The R package attached to LFQbench project merged the six identification results, visualized the distribution of the relative quantitative and the experimental ratios (Fig. 5a) and reported the number of peptides and proteins fell within the valid ratios (which were defined as a range of five standard deviations from the average ratio by LFQbench, Fig. 5b). As we can see, on the one hand, Alpha-XIC significantly increased the number of peptides and proteins fell within valid ratios compared to OpenSWATH (14.8% and 12.1% improvements for human peptides and proteins; 8.3% and 6.1% for yeast peptides and proteins; 16.6% and 13.8% improvements for *E. coli* peptides and proteins), on the other hand, the results of Alpha-XIC slightly enlarged the median deviations between the experimental ratios and the expected ratios (Alpha-XIC introduced the median deviations: 0.00 and 0.00 for human peptides and proteins, respectively, −0.02 and −0.02 for yeast peptides and proteins, respectively, 0.17 and 0.15 for *E. coli* peptides and proteins, respectively, compared to the median deviations by OpenSWATH: 0.00 and 0.00 for human peptides and proteins, respectively, −0.01 and 0.00 for yeast peptides and proteins, respectively, 0.14 and 0.10 for *E. coli* peptides and proteins, respectively). In other words, unlike Avant-garde tool (Jacome *et al*., 2020) which achieved fewer identification but more accurate global quantification, we believed Alpha-XIC delivered more identification at the expense of the reduction of integral quantitative accuracy. This may be due to the fact that Alpha-XIC recovered some of the interfered peptides, but OpenSWATH’s quantitative algorithm cannot eliminate the influence of the interference. A better quantification algorithm for peptides with interference correction needs to be developed in the future.

**Fig. 5.**
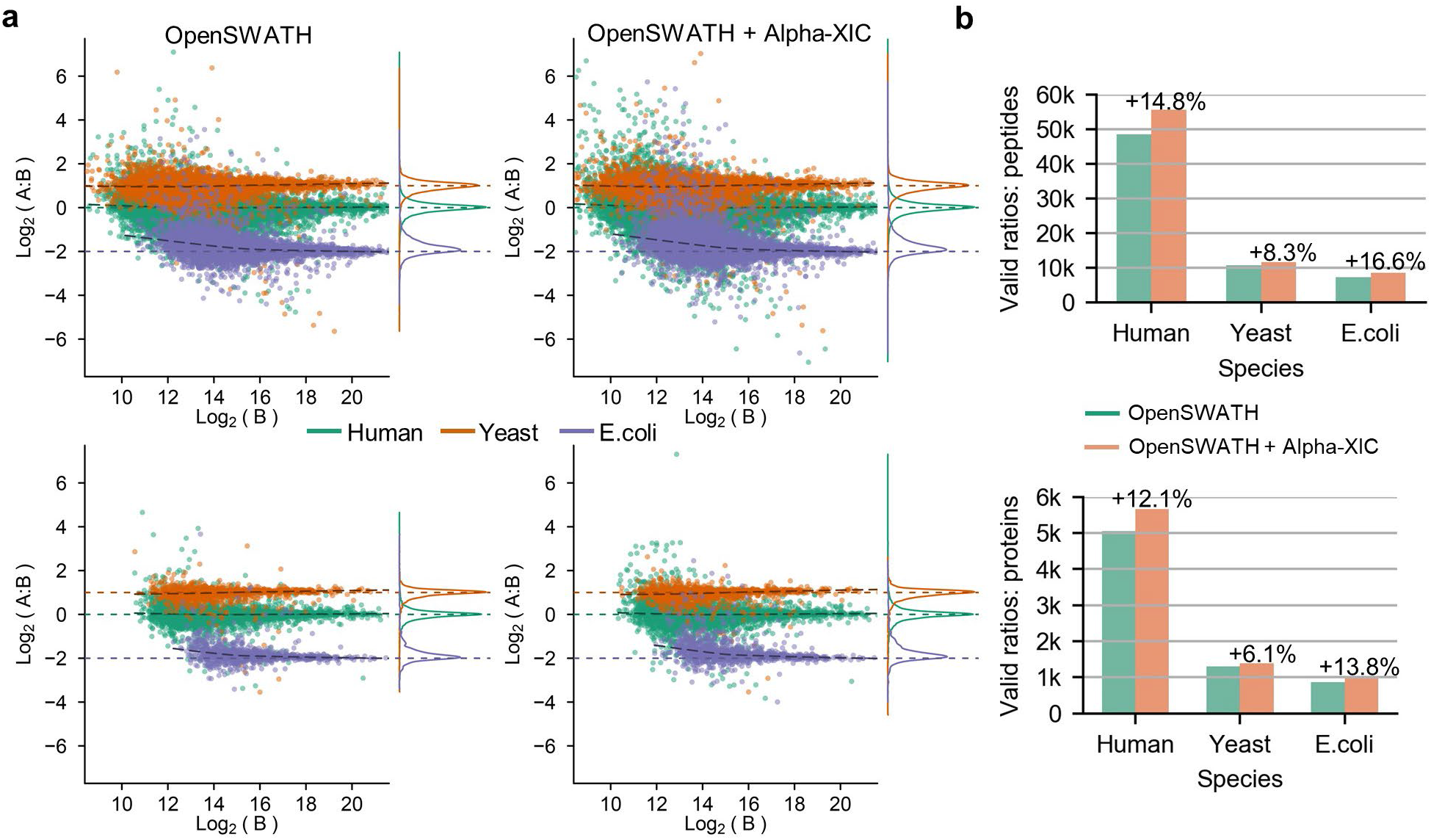
Performance evaluation of Alpha-XIC on LFQbench dataset. **a)** Visualization of the distribution of the relative quantitative and the experimental ratios using the LFQbench R package. **b)** Comparisons of the numbers of valid ratios on the peptide and protein level.

## 4 Conclusion

In this study, we propose a neural network, Alpha-XIC, to score the coelution of peak groups.

Instead of manually designing functions, Alpha-XIC characterizes the coelution by learning the peak group itself, which makes it possible to eliminate the influence of interference and achieve the robust coelution score. We found that Alpha-XIC could give a high coelution score to a peak group even in the circumstance of severe interference. By appending the coelution scores from Alpha-XIC to the initial scores from OpenSWATH, Alpha-XIC obtained improvements by 16.7% ~ 49.1% in terms of the number of identified precursors at 1% FDR for HeLa dataset. Besides, on the LFQbench dataset, Alpha-XIC increased the number of peptides and proteins fell within the valid ratios range by up to 16.6% and 13.8%, respectively, compared to the identification by OpenSWATH solely.

As models like Alpha-Frag (Song *et al*., 2021) have implemented the presence prediction of fragment ions of a peptide, in the future, Alpha-XIC can extend the ions involved in the construction of peak groups to precursor ions, unfragmented precursor ions and the predicted present fragment ions as well as their isotopic ions, rather than just the few fragment ions provided in the spectral library. In this way, Alpha-XIC takes into account the coelution of all ions derived from one targeted peptide, thus making full use of the coelution information and improving the identification of DIA data further.

